# Myeloid cell-associated resistance to PD-1/PD-L1 blockade in urothelial cancer revealed through bulk and single-cell RNA sequencing

**DOI:** 10.1101/2020.09.16.300111

**Authors:** Li Wang, John P. Sfakianos, Kristin G. Beaumont, Guray Akturk, Amir Horowitz, Robert Sebra, Adam M. Farkas, Sacha Gnjatic, Austin Hake, Sudeh Izadmehr, Peter Wiklund, William K Oh, Peter Szabo, Megan Wind-Rotolo, Kezi Unsal-Kacmaz, Xin Yao, Eric Schadt, Padmanee Sharma, Nina Bhardwaj, Jun Zhu, Matthew D. Galsky

## Abstract

Adaptive immunity and tumor-promoting inflammation exist in delicate balance in individual tumor microenvironments; however, the role of this balance in defining sensitivity and resistance to PD-1/PD-L1 blockade therapy in urothelial cancer and other malignancies is poorly understood. We pursued an unbiased systems biology approach using bulk RNA sequencing data to examine pre-treatment molecular features associated with sensitivity to PD-1/PD-L1 blockade in patients with metastatic urothelial cancer and identified an adaptive_immune_response module associated with response and an inflammatory_response module and stromal module associated with resistance. We mapped these gene modules onto single-cell RNA sequencing data demonstrating the adaptive_immune_response module emanated predominantly from T, NK, and B cells, the inflammatory_response module from monocytes/macrophages, and the stromal module from fibroblasts. The adaptive_immune_response:inflammatory_response module expression ratio in individual tumors, reflecting the balance between antitumor immunity and tumor-associated inflammation and coined the 2IR score, best correlated with clinical outcomes and was validated in an independent cohort. Individual monocytes/macrophages with low 2IR scores demonstrated upregulation of proinflammatory genes including IL1B and downregulation of antigen presentation genes, were unrelated to classical M1 versus M2 polarization, and were enriched in pre-treatment peripheral blood from patients with PD-L1 blockade-resistant metastatic urothelial cancer.

**Single sentence summary:** Proinflammatory monocytes/macrophages, present in tumor and blood, are associated with resistance to immune checkpoint blockade in urothelial cancer.

## Introduction

Standard treatment for metastatic urothelial cancer (UC) of the bladder has historically been limited to platinum-based chemotherapy. However, the treatment landscape has recently experienced a major shift with the introduction of several PD-1/PD-L1 immune checkpoint inhibitors (CPI) into the armamentarium.(*1–5*) These therapies are characterized by durable responses, often measured in years, but achieved in only a subset of ∼15-25% of patients. This therapeutic profile has led to intensive investigation into mechanisms of intrinsic resistance in pursuit of biomarkers and combination strategies to extend the benefits of CPI to additional patients.

Responses to CPI are thought to be predicated on a pre-existing anti-tumor T cell response restrained due to adaptive immune resistance.(*6*) Indeed, measures of T cell infiltration, IFNγ-related gene signatures, and PD-L1 expression, colloquially referred to as reflecting “hot tumors”, have all been correlated with response to CPI in patients with UC.(*1, 3*) These biomarkers are also generally positively correlated with one another, likely reflecting largely redundant biology.(*7*) While PD-L1 expression is the only biomarker integrated into clinical practice to inform CPI treatment in UC to date, PD-L1 testing alone conveys modest predictive information. Several groups have shown that higher tumor mutational burden (TMB) is associated with an increased likelihood of response to CPI independent of measures of adaptive immune resistance.(*7–9*) These findings suggest that combining multiple cancer cell- and tumor microenvironment (TME)- related features may be required to engender mechanistic insights and refine clinical decision-making.

Tumor-promoting “chronic” inflammation, now recognized as a hallmark of cancer pathogenesis, involves a TME shaped by activated fibroblasts, endothelial cells and innate immune cells.(*10*) Monocytes and macrophages, key participants in tumor-promoting inflammation, have been implicated in resistance to various cancer therapies.(*11, 12*) Remarkably, despite the widely held notion that subsets of myeloid cells in the TME impair T cell immunity, there have been relatively few studies involving human specimens linking a myeloid-inflamed TME with CPI resistance.(*13*) Further, the TME is comprised of diverse cell types and/or states orchestrating intricate interactions. Which of these various cellular populations and interactions underlie dominant mechanisms of CPI resistance is poorly understood; yet, such a holistic understanding is critical to prioritizing putative biomarkers and therapeutic targets for further investigation.

To address the aforementioned knowledge gaps, we pursued an unbiased systems biology approach using bulk RNA sequencing data to examine pre-treatment molecular features associated with CPI outcomes in metastatic UC and identified three key gene modules: 1) an adaptive_immune_response module enriched in adaptive immune response genes and associated with longer survival, 2) an inflammatory_response module enriched in inflammation and innate immune genes and associated with shorter survival, and 3) a stromal module enriched in epithelial mesenchymal transition (EMT)- and extracellular matrix (ECM)-related genes and associated with shorter survival. The adaptive_immune_response:inflammatory_response module expression ratio, coined the *2IR score*, had the largest effect on clinical outcomes and was validated in an independent cohort. We then used single-cell RNA sequencing (scRNA-seq) data generated from UC bladder specimens to uncover the cellular composition underlying the three gene modules demonstrating that the adaptive_immune_response module emanated predominantly from T, NK, and B cells, the inflammatory_response module from monocytes/macrophages, and the stromal module from cancer associated fibroblasts (CAFs). Individual monocytes/macrophages with low 2IR scores demonstrated upregulation of proinflammatory genes including *IL1B* and downregulation of antigen presentation genes, were unrelated to classical M1 versus M2 polarization, and were enriched in the pretreatment blood of patients with CPI-resistant metastatic UC. Thus, the balance of adaptive immunity and tumor-associated inflammation drives outcomes with CPI in UC and resistance associated with the latter may be mediated by a population of proinflammatory monocytes/macrophages detectable in both the TME and peripheral blood.

## Results

### An adaptive_immune_response, inflammatory_response, and stromal gene module were associated with CPI outcomes in patients with UC

To identify molecular features associated with survival in CPI-treated patients with metastatic UC, we utilized bulk RNA sequencing and TMB data from the IMvigor 210 study, a large single arm phase 2 trial testing the PD-L1 inhibitor, atezolizumab (Figure 1a).(*1, 7, 14*) This cohort has been previously described and additional details are provided in Table S1; RNA sequencing and TMB data were available for 348 and 272 patients, respectively.(*7*) We pursued step-wise identification of consistently co-expressed gene modules, which focused on identifying gene modules associated with overall survival (OS) and utilizing gene modularity to enrich for true signals (Figure 1b, Figure S1; see Methods). Given the correlation between TMB and response to CPI in UC(*7, 15*), we explored genes associated with OS conditioning on TMB (see Methods) and identified a signature consisting of 1193 genes associated with longer OS. To refine this signature, we performed meta-analysis of co-expression patterns(*16, 17*) using both the IMvigor 210 and The Cancer Genome Atlas (TCGA) UC datasets and identified a consistently co-expressed gene module comprised of 483 genes (see Methods and Extended Data). Gene set enrichment analysis revealed that this module was highly enriched in adaptive immune response-related genes (Figure 1c and Figure S1) and was therefore labeled the adaptive_immune_response module.

**Figure 1.**
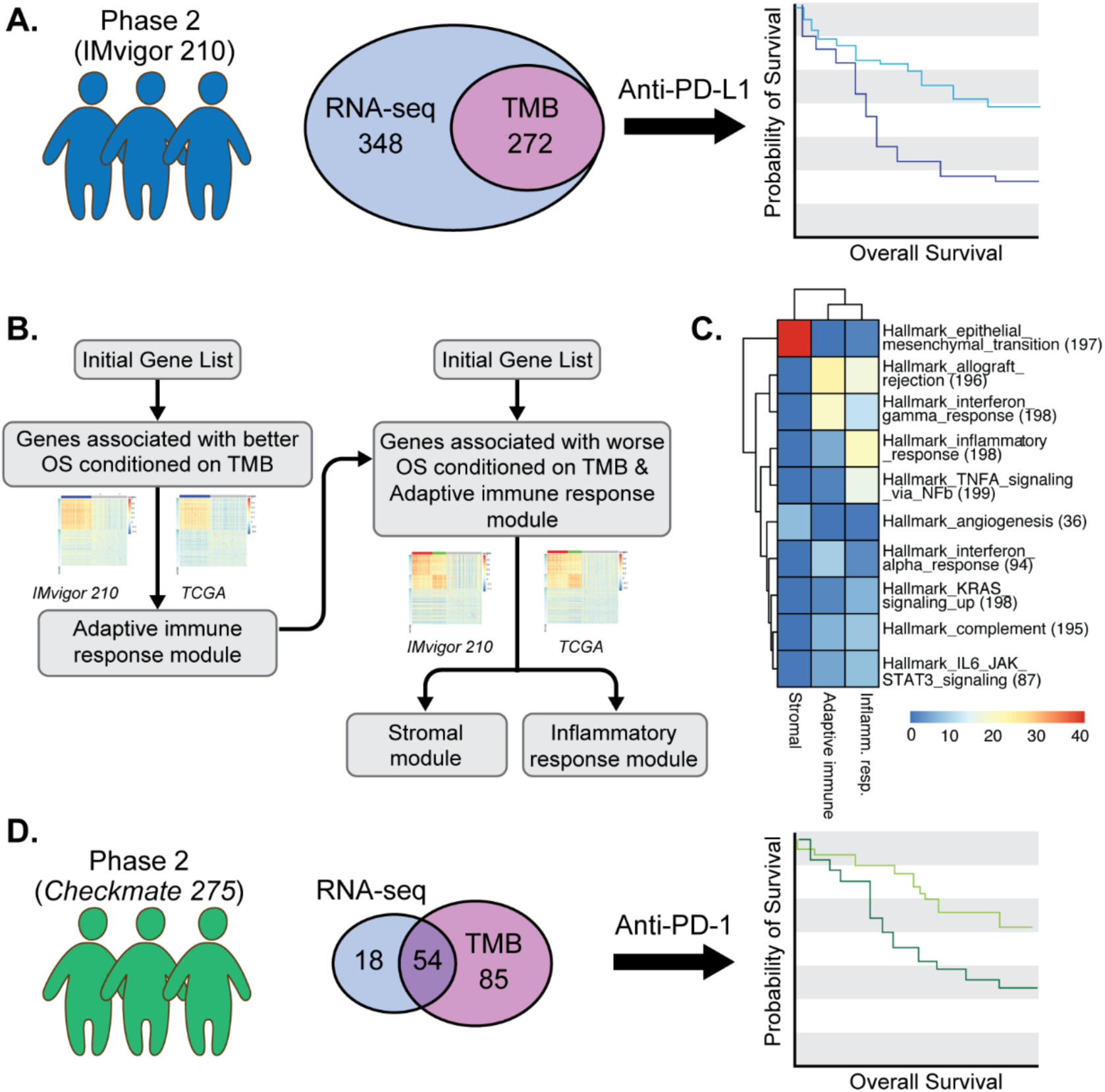
Cohorts and workflow for discovery of gene modules associated with sensitivity and resistance to anti-PD-1/PD-L1 treatment in metastatic urothelial cancer. A. IMvigor 210 was a single-arm phase 2 study investigating PD-L1 inhibition with atezolizumab in patients with metastatic urothelial cancer. The illustration depicts the numbers of patients with available pre-PD-L1 inhibition RNA-sequencing (RNA-seq) data, tumor mutational burden (TMB) data, or both, derived from archival tumor specimens available for the current analysis. B. Step-wise approach to the identification of consistently co-expressed gene modules, conditioned on TMB, associated with better overall survival or worse overall survival with PD-L1 blockade treatment in patients with metastatic urothelial cancer. Data from The Cancer Genome Atlas (TCGA) urothelial bladder cancer dataset was used to identify consistently co-expressed gene modules (see Methods). C. Hallmark pathways enriched in the adaptive_immune_response, inflammatory_response, and stromal gene modules using Fisher’s exact test (nominal two-sided p-value <1e-5). Color corresponds to the –log10 of the p-value. D. Checkmate 275 was a single-arm phase 2 study investigating PD-1 inhibition with nivolumab in patients with metastatic urothelial cancer. The illustration depicts the number of patients with available pre-PD-1 inhibition RNA-sequencing data, TMB data, or both derived from archival tumor specimens used for validation of the association between the adaptive_immune_response, inflammatory_response, and stromal gene modules and outcomes with PD-1/PD-L1 blockade in metastatic urothelial cancer.

In the second step, we further analyzed the IMvigor 210 dataset to identify genes associated with survival conditioning on both TMB and the adaptive_immune_response module (Figure 1b). We identified 1498 genes associated with shorter OS. We again applied meta-analysis of co-expression patterns(*18, 19*) in the IMvigor 210 and TCGA UC datasets and identified two consistently co-expressed gene modules for further analysis. The first module associated with shorter OS, consisting of 437 genes, was enriched in inflammation and innate immune genes (Figure 1c, Figure S1, and Extended Data), and was labeled the inflammatory_response module. The second module associated with shorter OS, consisting of 287 genes, was enriched in epithelial mesenchymal transition (EMT)- and extracellular matrix (ECM)-related genes (Figure 1c and Extended Data) and consistent with our prior work(*20*) was named the stromal module. Importantly, expression of the inflammatory_response and stromal modules were both positively correlated with the adaptive_immune_response module (Figure S2) such that their disparate impact on OS was only revealed using our stepwise approach (Figure S3) and suggesting that the balance of these features in individual tumors may impact CPI outcomes.

We next sought to define the independent contribution of the three gene modules to outcomes with CPI in the IMvigor 210 cohort. Models combining both the adaptive_immune_response and inflammatory_response modules (see Methods), particularly the adaptive_**I**mmune_response:**I**nflammatory_response module expression **R**atio (hereafter referred to as the **2IR score**), demonstrated the largest effect size on OS and similar findings were observed with objective response rate (Figure 2a-c and Table S2). Importantly, when both the inflammatory_response and stromal modules were entered into a multivariable model along with the adaptive_immune_response module and TMB, the stromal module was no longer independently associated with OS (Figure 2a and Table S2). These findings indicated that (a) the balance of cellular and molecular events underlying the adaptive_immune_response versus inflammatory_response modules within an individual UC TME may dictate outcomes with CPI and (b) the negative impact of the stromal module on outcomes may be largely indirect and mediated via the events underlying the inflammatory_response module (Figure 2a).

**Figure 2.**
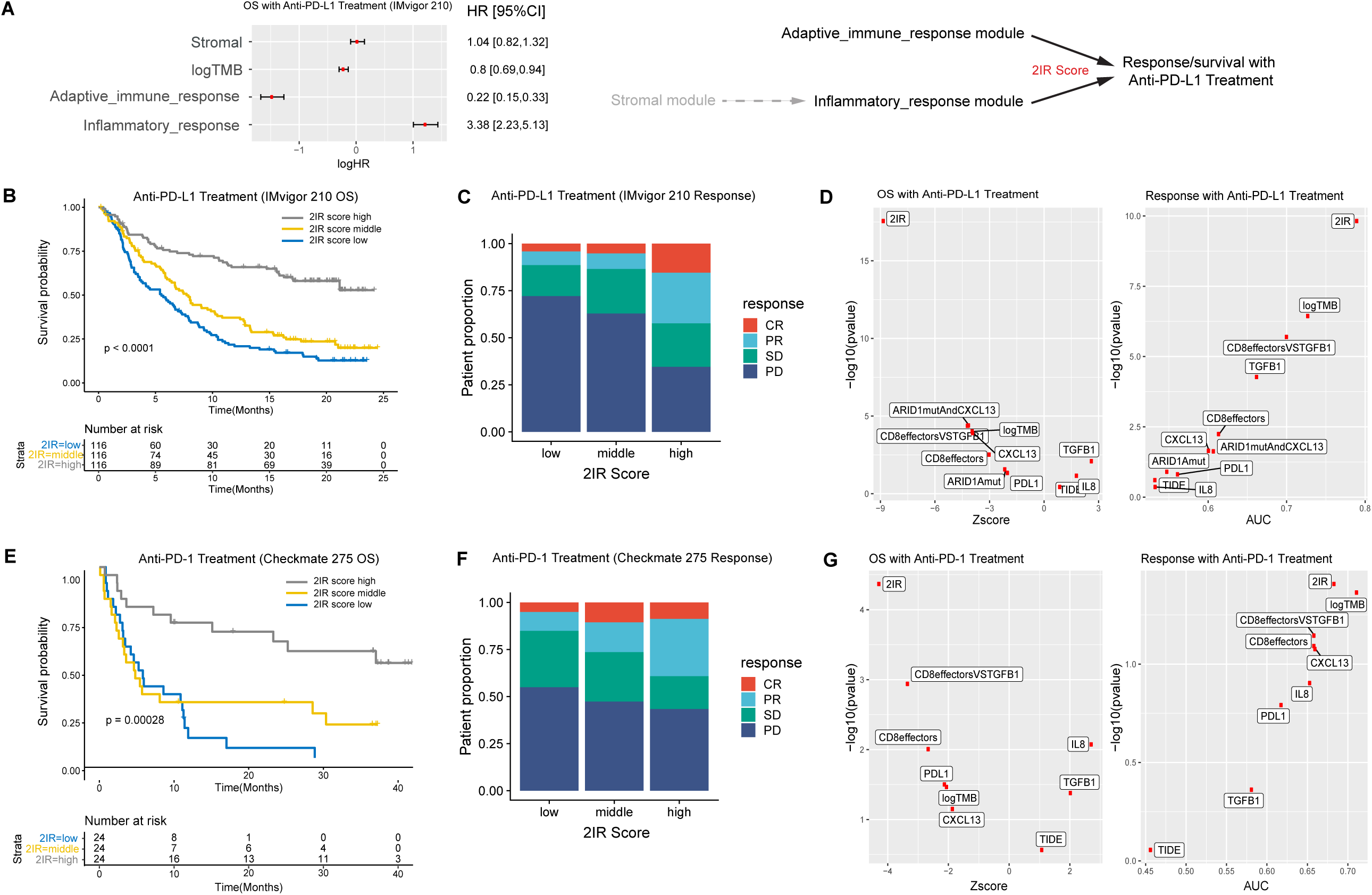
The adaptive_immune_response and inflammatory_response gene modules, and the ratio of module expression termed the 2IR score, are associated with clinical outcomes with PD-1/PD-L1 blockade in patients with metastatic urothelial cancer. A. Multivariable Cox regression model for overall survival (OS; n=272 patients with RNA sequencing and tumor mutational burden (TMB) data) including adaptive_immune response, inflammatory_response, and stromal module expression, as well as TMB from the IMvigor 210 cohort (HR, hazard ratio; 95% CI, 95% confidence interval; error bars represent 95% CI of the HRs). Module expression and TMB were standardized before entering the Cox regression model. The plot indicates log HRs while annotation provides HRs. Schematic representation of the relationship of the adaptive_immune_response, inflammatory_response, and stromal modules and outcomes with atezolizumab indicating potential indirect role of the stromal module on resistance mediated more directly through the inflammatory response module and the **2IR score** representing the adaptive_**I**mmune response:**I**nflammatory_response module expression **R**atio. B. Kaplan-Meier curve for overall survival (OS) stratified by the 2IR score cut at tertiles in the IMvigor 210 cohort (n=348 patients with RNA sequencing data; log-rank p value shown). C. Objective response rate with PD-L1 blockade in the IMvigor 210 cohort according to the 2IR score (cut at tertiles). For each 2IR score tertile, bar graphs depict the percentage of patients achieving a complete response (CR), partial response (PR), stable disease (SD), or progressive disease (PD) as the best objective response with PD-L1 blockade. D. The association between each biomarker (or biomarker combination) and overall survival (OS) in the IMvigor 210 cohort was evaluated using the Z-score by univariate Cox regression analysis and the p-value by log likelihood ratio test (left). The association between each biomarker and response to PD-L1 blockade (CR/PR versus SD/PD) was evaluated using the area under curve (AUC) score and the p-value by the Wald’s test in univariate logistic regression (right). E. Kaplan-Meier curves for overall survival (OS) stratified by the 2IR score (cut at tertiles) in the Checkmate 275 cohort (n=72 patients with RNA sequencing data; log rank p value shown). F. Objective response rate with PD-1 blockade in the Checkmate 275 cohort according to the 2IR score (cut at tertiles). For each 2IR score tertile, bar graphs depict the percentage of patients achieving a complete response (CR), partial response (PR), stable disease (SD), or progressive disease (PD) as the best objective response with PD-1 blockade. G. The association between each biomarker (or biomarker combination) and overall survival (OS) in the Checkmate 275 cohort was evaluated using the Z-score by univariate Cox regression analysis and the p-value by log likelihood ratio test (left). The association between each biomarker and response to PD-1 blockade (CR/PR versus SD/PD) was evaluated using the area under curve (AUC) score and the p-value by the Wald’s test in univariate logistic regression (right).

Given the potential practical advantages of smaller sets of module genes for validation and clinical biomarker development, we identified the top-ranked genes within the three modules (see Methods, Table S3, and Figure S4). Module scores derived from these smaller gene sets demonstrated similar associations with OS compared with scores derived from the full gene sets (Table S4).

### The three gene modules were validated in an independent dataset of patients with metastatic UC treated with PD-1 blockade and conveyed information beyond previously identified features

For validation, we utilized TMB and RNA sequencing data from the Checkmate 275 study, a single-arm phase 2 trial evaluating the PD-1 inhibitor, nivolumab, in patients with metastatic UC (Figure 1d).(*3*) This cohort has been previously described, with further detail provided in Table S1; RNA sequencing and TMB data were available for 72 and 139 patients, respectively.(*3*) Gene module expression was defined using the top-ranked genes identified in the IMvigor 210 cohort. Within the limitations of the smaller sample size, adaptive_immune_response, inflammatory_response, and stromal module expression demonstrated similar associations with OS and response rate in the Checkmate 275 cohort (Table S5). As observed in the IMvigor 210 cohort, the 2IR score in the Checkmate 275 cohort demonstrated the largest effect on CPI outcomes (Figure 2e,f).

Our two cohorts involving CPI treatment were single arm clinical trials precluding a full understanding of the predictive versus prognostic nature of the gene modules. However, the effect of the 2IR score on OS in TCGA UC dataset, a cohort of patients with muscle-invasive UC of the bladder treated with curative-intent cystectomy, was less dramatic (Figure S5). Coupled with the correlation with CPI objective response rate in addition to OS, these findings reinforced that the 2IR score may impart predictive rather than solely prognostic information.

Other features associated with outcomes with CPI in UC and other cancers have been reported including PD-L1 protein expression, the tumor immune dysfunction and exclusion (TIDE) and CD8 effector T cell gene signatures, *ARID1A* mutation status, and *CXCL13*, *TGFB1, or CXCL8 (IL8)* gene expression (see Figure S6 for correlation between these features and the 2IR score).(*7, 21–23*) Importantly, the 2IR score was associated with favorable performance characteristics relative to these other measures in both the IMvigor 210 and Checkmate 275 cohorts (Figure 2d,g). Taken together, the 2IR score, representing the balance of expression of the adaptive_immune_response and inflammatory_response gene modules within individual TMEs, is associated with response rate and OS in CPI-treated patients and conveys information beyond that achieved with previously identified features.

### Lower 2IR scores, and stromal module expression, were associated with a paucity of T cells in cancer cell nests

To examine the TME of UC defined by the 3 gene modules, we employed a tissue profiling approach known as multiplexed immunohistochemical consecutive staining on a single slide (MICSSS)(*24, 25*) on a subset of 19 specimens from the Checkmate 275 cohort with available unstained slides and matched RNA sequencing data. MICSSS revealed that specimens with higher 2IR scores generally exhibited CD8+ T cell infiltration invading cancer cell nests and occasional tertiary lymphoid-like structures (Figure 3a,b and Figure S7) whereas specimens with lower 2IR scores demonstrated dense stroma (Figure 3 c,d). Hence, the 2IR score was associated with histologic features reminiscent of adaptive immunity versus tumor-associated inflammation.

**Figure 3.**
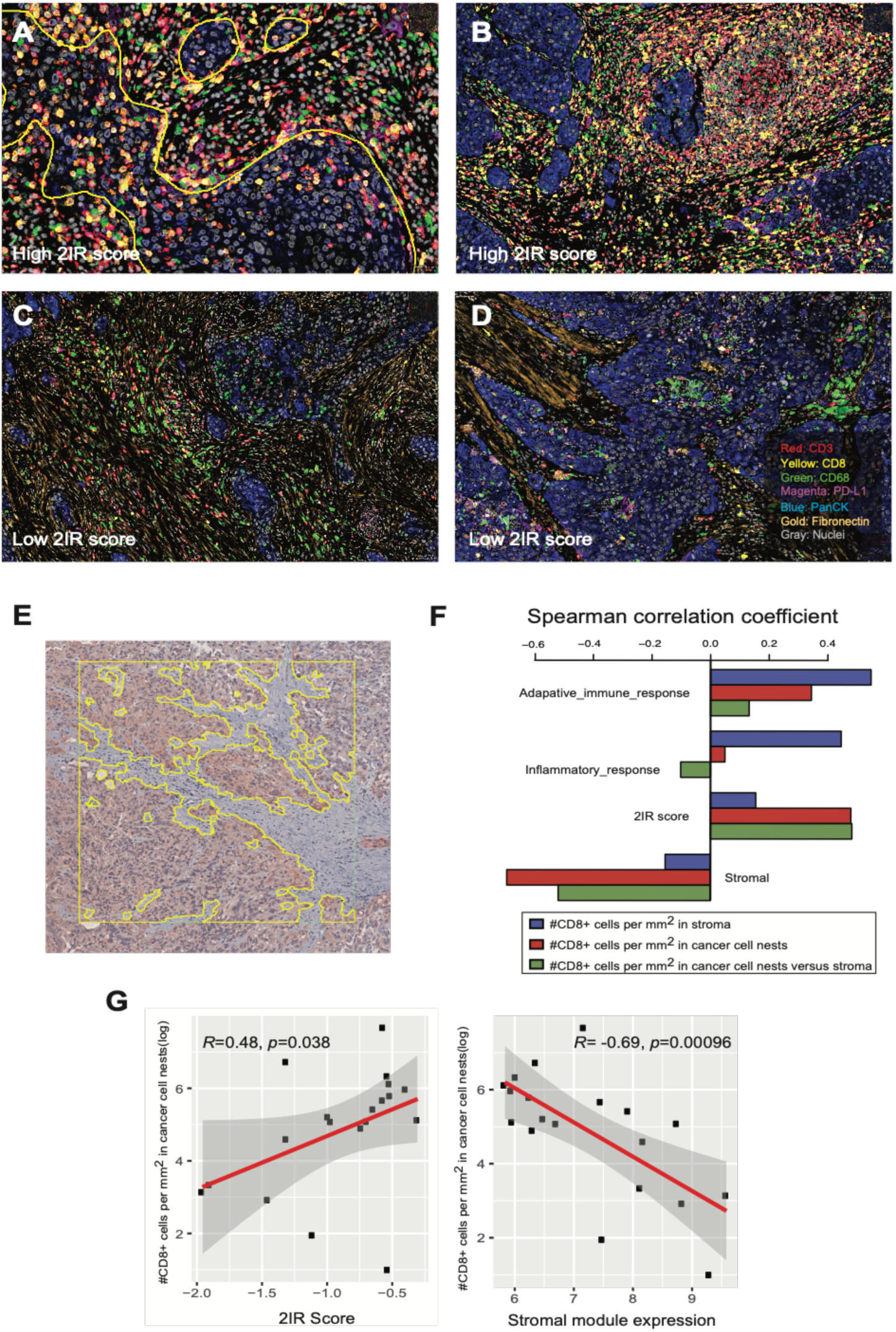
The gene modules are associated with spatial organization of immune cells in the tumor microenvironment. A-D. Representative images of multiplexed immunohistochemical consecutive staining on a single slide (MICSSS) demonstrating abundance of CD8+ T cells (A, B) and tertiary lymphoid-like structures (B) in specimens with high 2IR scores and a paucity of CD8+ T cells and prominent stroma (C, D) in specimens with a low 2IR scores. Yellow outline in panel A represents demarcation of cancer cell nests. All slides were initially scanned at 20x magnification. E. Representative image of urothelial cancer specimen demonstrating region of interest (ROI), designated by the square, and machine learning-based segmentation of cancer cell nest and stromal zones to define T cell localization in the tumor microenvironment using pancytokeratin immunohistochemical staining, designated by the yellow outline bordering cytokeratin-expressing cells. F. Spearman’s correlation between enumeration of CD8+ T cells localized to cancer cell nests or stromal zones and adaptive_immune_response module, inflammatory_response module, 2IR score, or stromal gene module expression. The results are based on analysis of 76 ROIs across 19 specimens with both immunohistochemistry and RNA sequencing data from the Checkmate 275 cohort. G. Correlation between enumeration of CD8+ T cells localized to cancer cell nests and the 2IR score (left) or stromal module expression (right). The results are based on analysis of 76 ROIs across 19 specimens with both immunohistochemistry and RNA sequencing data from the Checkmate 275 cohort. Spearman’s correlation was used to determine the correlation coefficient R and p value.

We, and others, previously showed that UC specimens with increased stroma-related gene expression were characterized by T cells spatially separated from cancer cell nests, commonly referred to as the “excluded” phenotype.(*7, 20*) Therefore, to quantify the localization of T cells according to gene module expression, we defined cancer cell and stromal zones based on pan-cytokeratin staining using a machine learning segmentation tool and examined CD8+ expression in 76 regions of interest across the 19 specimens (see Methods and Figure 3e). Lower 2IR scores, or higher stromal module expression, correlated with decreased CD8+ T cell enumeration in cancer cell nests (Figure 3f,g). These findings suggested that CPI resistance associated with lower 2IR scores may be related to impairment of T cell trafficking and/or function prompting us to further probe the cellular origins of gene module expression.

### The three gene modules emanated predominantly from distinct cellular components of the TME as identified by single cell RNA sequencing

Our gene modules were derived from bulk RNA sequencing data from archival UC specimens obtained pre-treatment with CPI, the vast majority of which represented invasive primary tumors (Table S1). Therefore, to map the cellular origins of the three gene modules (Figure 4a), we performed droplet-based encapsulation single cell RNA sequencing (scRNA-seq) on an analogous set of eight freshly resected invasive UC bladder specimens as well as two specimens derived from adjacent grossly normal urothelium using the 10x Genomics Chromium system (see Methods). The characteristics of the cohort are detailed in Table S6. After excluding cells not passing quality control (see Figure S8 for QC plots), 19,708 cells from the 10 samples were analyzed. A median of 1456 genes were detected per cell. We performed graph-based clustering as implemented in the *Seurat* package.(*26*) A two-stage clustering approach was employed in which cells were first grouped into major clusters and subsequently further partitioned into minor clusters (see Methods). Canonical marker genes revealed nine major cell populations including T- and NK cells, B-cells, myeloid-lineage cells, non-hematopoietic stromal cells, and epithelial cells (Figure 4b,c). Granulocyte populations were not well captured by 10x Chromium scRNA-seq as has been noted in prior studies.(*27*)

**Figure 4.**
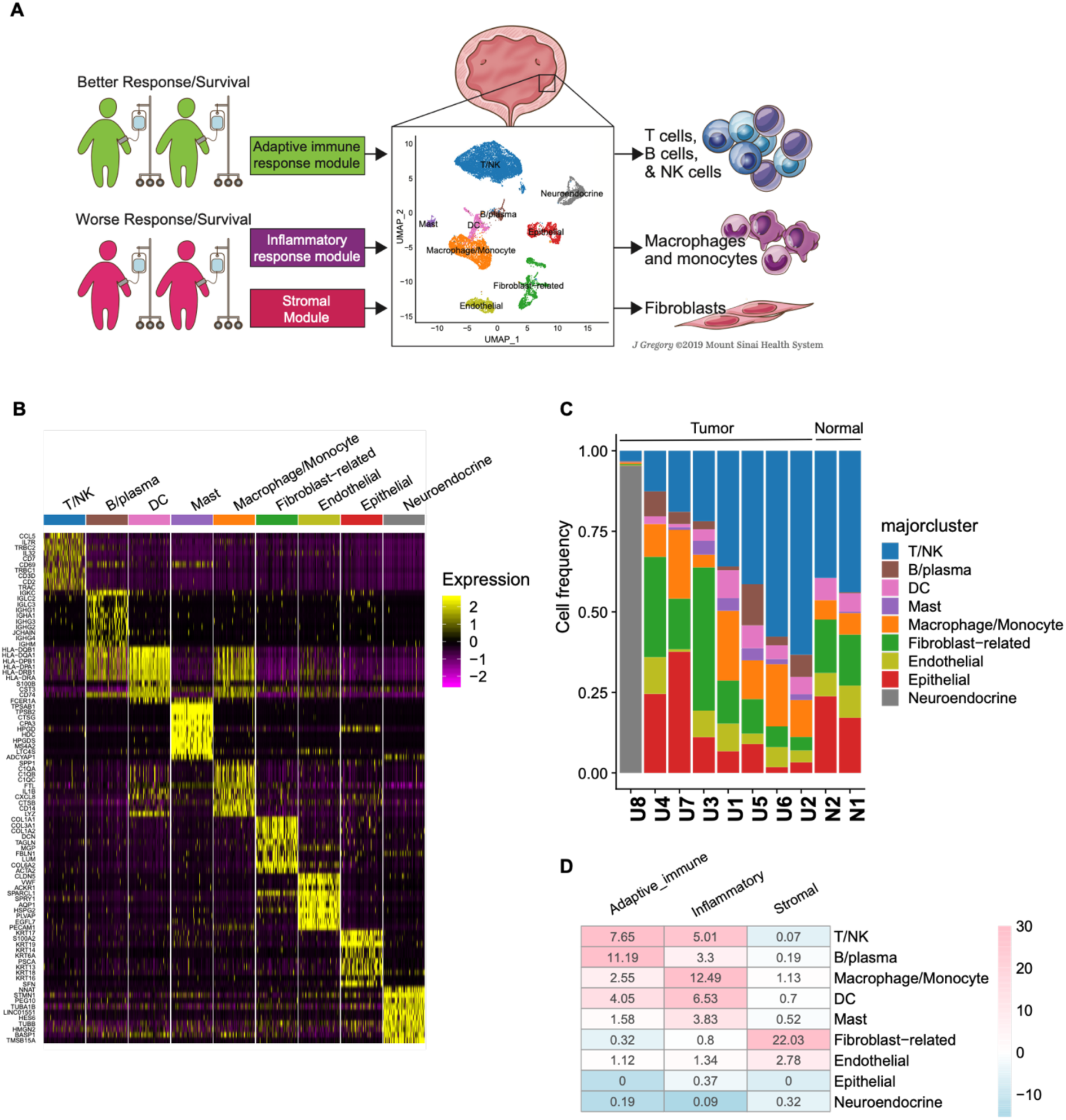
Defining the cellular origins of adaptive_immune_response, inflammatory_response, and stromal module gene expression using single-cell RNA sequencing. A. Schematic representation of projection of gene modules identified using bulk RNA sequencing data linked to outcomes with anti-PD-1/PD-L1 treatment onto single-cell RNA sequencing data generated from a separate cohort of invasive urothelial bladder cancer specimens. The illustration depicts nine major cell clusters visualized using Uniform Manifold Approximation and Projection (UMAP) across eight urothelial cancer specimens and two adjacent normal urothelial cancer specimens profiled using droplet-based encapsulation single-cell RNA sequencing. The adaptive_immune_response, inflammatory_response, and stromal modules identified using bulk RNA sequencing data from clinical trial cohorts were projected onto the single-cell RNA sequencing data to define the predominant cellular sources of the respective module gene expression. B. Single-cell expression of top 10 overexpressed genes in each major cell cluster. Heatmap visualization color-coding the scaled gene expression level for selected marker genes (rows). Visualized are 500 randomly selected cells per cluster. C. Frequency of cell populations in individual samples included in the single-cell RNA sequencing cohort. For each sample, bar graphs depict the percentage of cells in clusters associated with each population. Samples were ranked according to T/NK cell frequency. Normal indicates samples obtained for urothelial tissue that was considered grossly normal by visual inspection adjacent to site of harvested tumor tissue. D. Heatmap of overlap between genes comprising the adaptive_immune_response, inflammatory_response, and stromal modules and genes overexpressed in each of the major cell clusters in the single-cell RNA sequencing cohort. The number in each cell corresponds to the odds ratio for the corresponding overlap between genes, the color corresponds to the –log10 p-value (for enrichment) or log10 p value (for depletion) by two-sided Fisher’s exact test.

To determine the predominant origins of the adaptive_immune response, inflammatory_response, and stromal modules, the expression pattern of the module genes was assessed among the major cell populations (see Methods). The adaptive_immune_response module was enriched among genes overexpressed by T, NK, and B cells, the inflammatory_response module among genes overexpressed by monocytes/macrophages, and the stromal module among genes overexpressed by fibroblasts (Figure 4a,d).

Partitioning of the major cell clusters through a second round of analysis (see Methods) revealed 50 minor cell clusters (Figures 5a). This high-resolution characterization of UC specimens demonstrated that while the gene modules were expressed predominantly by certain cell types, diversity in module expression between and among cell types was observed suggesting that interplay among different cell types in the TME contributed to module expression (Figure 5a). Given their key role in contributing to the inflammatory_response module, the monocyte/macrophage clusters are described further herein while the remainder of the minor cell clusters are described in the Supplemental Results and Figures S9-15.

**Figure 5.**
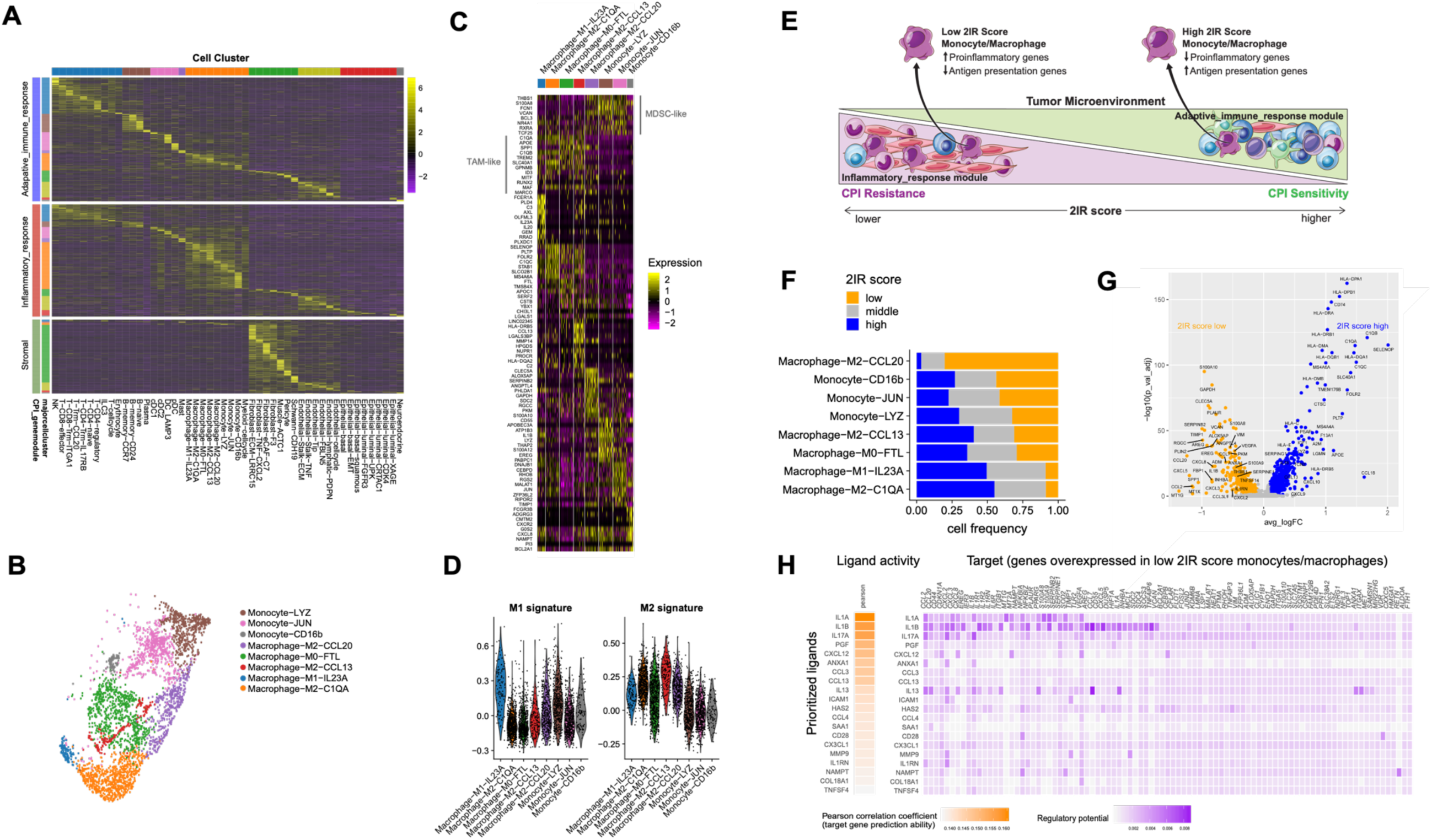
The inflammatory_response module is traced predominantly to monocytes/macrophages and low 2IR score monocytes/macrophages are characterized by increased expression of proinflammatory genes and decreased expression of antigen presentation genes. A. Heatmap visualizing the expression of adaptive_immune_response, inflammatory_response, and stromal module genes across each of the major and minor cell clusters, B. Eight minor monocyte/macrophage cell clusters visualized using Uniform Manifold Approximation and Projection (UMAP) across eight urothelial cancer specimens and two adjacent normal urothelial cancer specimens profiled using droplet-based encapsulation single-cell RNA sequencing. C. Monocyte/macrophage cell populations in the single-cell RNA sequencing cohort. Heatmap visualization color-coding the scaled gene expression level for selected marker genes (rows). Visualized are 200 randomly selected cells per cluster or all cells when the cell cluster contained <200 cells. D. Expression level of M1 and M2 macrophage polarization signature genes in the monocyte/macrophage populations as assessed by the AddModuleScore() function in the Seurat package. E. Schematic representation of the relationship between the 2IR score in the urothelial cancer tumor microenvironment based on bulk RNA sequencing and the 2IR score in individual monocytes/macrophages based on single cell RNA sequencing. F. The frequency of cells with low, intermediate and high 2IR score within each monocyte/macrophage minor population. G. Volcano plot of genes differentially expressed between monocytes /macrophages with high versus low 2IR scores. P-value was calculated by Wilcoxon rank-sum test and then adjusted by Bonferroni correction. Genes with log fold change (FC) >0.1 and adjusted p-value <0.05 were considered as significant. H. Top-ranking ligands inferred to regulate genes upregulated in low 2IR score monocytes/macrophages according to NicheNet. Heatmap visualization of ligand activity and downstream target genes inferred to be regulated by each respective ligand.

We identified eight monocyte/macrophage clusters (Figure 5a-c). While some clusters expressed higher expression of M1 versus M2 signature genes (Figure 5d), or vice versa, heterogeneity of monocyte/macrophage subsets was observed beyond classical M1 and M2 polarization as has been documented in several analyses.(*28, 29*) The macrophage clusters resembled previously described “TAM-like macrophages” with increased expression of *APOE*, *C1QA*, *C1QB*, *SLC40A1* and *TREM2*.(*28, 30*) Two clusters demonstrated higher expression of *S100A* family genes, but lower M1 and M2 signature gene expression, and were annotated as monocyte-Jun and monocyte-LYZ. These clusters resembled previously described “MDSC-like macrophages” with overexpression of *THBS1*, *S100A8*, *FCN1* and *VCAN*.(*28*)

Our single-cell analysis reinforced that the cellular events underlying adaptive_immune_response module expression were characteristic of adaptive immunity while the cellular events underlying the inflammatory_response and stromal modules were characteristic of tumor-associated inflammation (Figure 5a). Coupled with our bulk gene expression data demonstrating that (a) the 2IR score best discriminated outcomes with CPI and (b) the stromal module did not convey independent information, these findings reinforced that the balance adaptive immunity and tumor-associated inflammation in an individual TME may dictate outcomes with CPI and that subsets of monocytes/macrophages may be key drivers of resistance associated with the latter.

### Monocytes/macrophages with low 2IR scores demonstrated increased expression of proinflammatory genes and decreased expression of antigen presentation genes

We next sought to better characterize the cellular state of monocytes/macrophages that might be linked to CPI resistance. Monocytes/macrophages are highly plastic, educated by cellular and signaling interactions in the TME, and play diverse roles in promoting and restraining anticancer immunity.(*31*) Despite inflammatory_response module expression emanating predominantly from monocytes/macrophages in the UC TME, our single cell characterization of UC specimens revealed diverse expression of the inflammatory_response module and adaptive_immune response module across individual macrophages/monocytes (Figure 5a and S16). Extending the concept of the balance of adaptive immunity and tumor-associated inflammation in UC TMEs to the single cell level, we calculated 2IR scores for each individual monocyte/macrophage identifying cells with low, intermediate, and high scores (Figure 5e, Figure S16, see Methods for details). While low 2IR score cells were observed across all monocyte/macrophage minor clusters, and were unrelated to M1 versus M2 classification, these cells were highly enriched in the Macrophage-M2-CCL2 cluster (OR=11.0, p-value<1e-16 by fisher’s exact test) and depleted in the Macrophage-M2-C1QA cluster (OR=0.14, p-value<1e-16 by fisher’s exact test; Figure 5f). Notably, the Macrophage-M2-CCL2 cluster demonstrated high expression of both “MDSC-like” and “TAM-like” signature genes (Figure 5d), suggesting that this population may represent an intermediate state between “MDSC-like” and “TAM-like” cells. In two UC specimens with matched scRNA-seq data from both tumor and adjacent normal tissue, the Macrophage-M2-CCL2 subset was absent from the adjacent normal specimens (N1 and N2) while present in the corresponding tumor specimens (6.2% and 6.7% for U1 and U2, respectively; p-value =0.046 by Fisher’s exact test; Figure S17).

Differential gene expression and gene set enrichment analysis of monocytes/macrophages with low versus high 2IR scores revealed significant upregulation of inflammatory pathways and top-ranking genes such as *IL1B*, *CXCL8 (IL8), SPP1*, and *CCL20* in the former while the latter demonstrated significant upregulation of genes and pathways related to antigen presentation and chemokines such as *CXCL9* and *CXCL10* (Figure 5g and S18).

Monocytes derived from the peripheral blood of patients with renal cancer have been previously shown to express proinflammatory cytokines and chemokines, including *IL1B, TNF, CCL20,* and *CXCL8 (IL8),* through an IL-1β-dependent mechanism.(*32*) We sought to define putative therapeutic targets implicated in polarizing monocytes/macrophages with low 2IR scores and not restrict our analysis to genes overexpressed in low 2IR score cells but rather seek upstream ligands implicated in driving the expression of such genes. We therefore used NicheNet (*33*), an approach that predicts ligands that modulate target gene expression by leveraging prior knowledge of signaling pathways and transcriptional regulatory networks (see Methods). Indeed, this analysis revealed that IL-1α and IL-1β were the top-ranked ligands inferred to regulate genes overexpressed in low 2IR score monocytes/macrophages (Figure 5h). Both IL1A and IL1B were predominantly expressed by monocytes/macrophages in our single cell cohort (Figure S19).

Thus, the balance of adaptive_immune_response and inflammatory_response module expression, as reflected by the 2IR score, extends to individual monocytes/macrophages within the TME (Figure 5e). Low 2IR score monocytes/macrophages, with upregulation of proinflammatory genes and downregulation of antigen presentation genes and unrelated to classical M1 versus M2 polarization, may play a key role in CPI resistance.

### Monocytes with low 2IR scores are enriched in the pre-treatment peripheral blood of patients with CPI-resistant metastatic UC

We next asked whether similar heterogeneity in 2IR scores was present in monocytes in the peripheral blood of patients with metastatic UC and whether these populations were associated with CPI resistance. Single-cell RNA sequencing data from peripheral blood mononuclear cells collected prior to the initiation of treatment with anti-PD-L1 CPI from five patients who achieved an objective response, and five patients who did not achieve an objective response, were utilized (see Methods and Figure S20). We calculated 2IR scores in individual monocytes identifying low, intermediate, and high 2IR score populations. Monocytes with low 2IR scores were significantly enriched in the peripheral blood of patients with CPI-resistant versus CPI-responsive metastatic UC (Figure 6a; p value =0.0048 by two-sided t-test). Alternatively, the five patients who responded to CPI could not be readily distinguished from the five patients with CPI resistant metastatic UC using monocyte minor subsets (Figure 6b) or individual genes (Figure 6c). Similar to our findings in the UC TME, low 2IR score monocytes in the pre-treatment peripheral blood of patients with metastatic UC demonstrated upregulation of proinflammatory genes and downregulation of antigen presentation genes (Figure 6d) and IL-1α and IL-1β were the top ranked ligands inferred to regulate this gene expression program. Therefore, low 2IR score proinflammatory monocytes/macrophages are present in both the TME and peripheral blood of patients with UC and are associated with CPI resistance.

**Figure 6.**
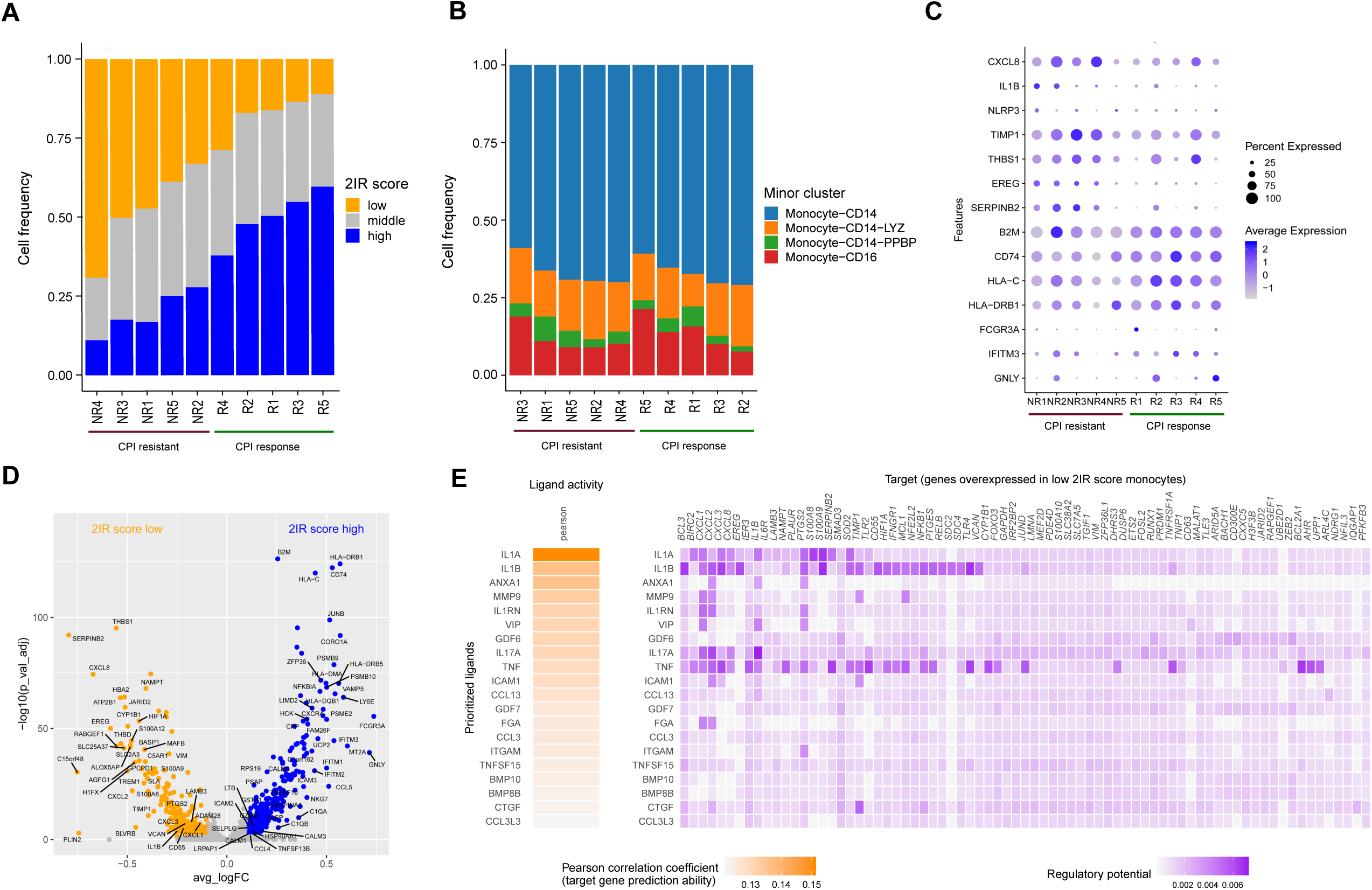
Low 2IR score monocytes are enriched in the pre-treatment peripheral blood of patients with metastatic urothelial cancer resistant to anti-PD-L1 treatment. Single-cell RNA sequencing data from peripheral blood mononuclear cells collected prior to the initiation of treatment from five patients with metastatic urothelial cancer who achieved an objective response, and five patients with metastatic urothelial cancer who did not achieve an objective response, to anti-PD-L1 immune checkpoint inhibition (CPI). A. The frequency of monocytes with low, intermediate and high 2IR scores in the pre-treatment peripheral blood of patients (n=10 patients) resistant or sensitive to anti-PD-L1 CPI. B. The frequency of monocyte minor cell populations in the pre-treatment peripheral blood of patients (n=10 patients) resistant or sensitive to anti-PD-L1 CPI. C. Dot plot of expression of select genes in monocytes from pre-treatment peripheral blood of patients (n=10 patients) resistant or sensitive to anti-PD-L1 CPI. D. Volcano plot of genes differentially expressed between peripheral blood monocytes with high and low 2IR score. P-value was calculated by Wilcoxon rank-sum test and then adjusted by Bonferroni correction. Genes with log fold change (FC) >0.1 and adjusted p-value <0.05 were considered as significant. E. Top-ranking ligands inferred to regulate genes upregulated in low 2IR score peripheral blood monocytes according to NicheNet. Heatmap visualization of ligand activity and downstream target genes inferred to be regulated by each respective ligand.

## Discussion

Tumor-associated inflammation is now recognized as a hallmark of cancer pathogenesis and major contributor to cancer treatment resistance.(*10, 34, 35*) However, tumor-associated inflammation and antitumor immunity coexist in delicate balance complicating dissecting the role of the former in mediating CPI resistance in studies using human specimens.(*13*) Using an unbiased systems biology approach, we demonstrated that an adaptive_immune_response gene module was associated with better CPI outcomes while an inflammatory_response gene module and stromal gene module were associated with worse CPI outcomes in patients with metastatic UC. We further demonstrated that: 1) the cellular origins of the modules were consistent with the adaptive_immune_response module reflecting key components of adaptive immunity (T and B-cells) and the inflammatory_response module and stromal modules reflecting key components of tumor-associated inflammation (monocyte/macrophages and activated CAFs, respectively), 2) expression of the three gene modules were highly correlated with one another, consistent with the coexistence and balance between antitumor immunity and tumor-associated inflammation, and the inflammatory_response and stromal modules were only discovered using our stepwise approach to module discovery, 3) the stromal module, linked to activated CAFs and remodeling of the extracellular matrix, did not convey independent information related to CPI outcomes beyond the inflammatory_response module suggesting a more indirect role (e.g., recruitment and education of myeloid cells), 4) the 2IR score, reflecting the balance of antitumor immunity and tumor-associated inflammation, best correlated with CPI outcomes, and 5) low 2IR score monocytes/macrophages were characterized by increased expression of proinflammatory genes and decreased expression of antigen presentation genes, could not be discerned based on classical M1 versus M2 polarization, and were enriched in the pre-treatment blood of patients with metastatic UC resistant to CPI. Together, our findings define a novel monocyte/macrophage population/cellular state associated with CPI resistance, highlight a potential approach to identify patients for therapies seeking to overcome monocyte/macrophage-related CPI-resistance, and delineate putative therapeutic targets to overcome such resistance.

Monocytes/macrophages have been linked to suppression of antitumor immunity across a range of malignancies via a variety of mechanisms.(*36–38*) Clinically tractable approaches to target myeloid cell-related immune suppression have remained elusive likely in part related to the plasticity of myeloid cells, redundancy of cytokines, and inability to identify patients for whom such strategies may be most appropriate. The NLRP3 inflammasome, implicated in the pathogenesis of several chronic inflammatory disorders, responds to a variety of stimuli leading to the generation of the active forms of the IL-1β and IL-18.(*39*) In our analysis, *NLRP3* was among the inflammatory_response module genes most highly associated with CPI resistance (Table S3) and *IL1B* was highly expressed in low 2IR score monocytes/macrophages. IL-1β was also among the top-ranked ligands inferred to regulate the low 2IR score monocyte/macrophage gene program in line with prior experimental data demonstrating that inflammatory cytokine and chemokine production from monocytes in patients with renal carcinoma was IL-1β-dependent.(*32*) IL-1β has been considered a “master regulator” of inflammation and involved in the tumor-promoting and immune suppressive function of myeloid cells.(*32, 40, 41*) IL-1β-deficient 4T1 mice were shown to harbor low levels of suppressive myeloid cells whereas in wild-type 4T1 mice, anti-IL-1β combined with anti-PD-1 therapy abrogated tumor growth.(*41*) Given the potential role of IL-1β in tumor promoting “chronic” inflammation, cancer incidence and mortality were analyzed in the Canakinumab Anti-inflammatory Thrombosis Outcomes Study (CANTOS) which randomized 10,061 patients with elevated serum C-reactive protein post-myocardial infarction to anti-IL-1β versus placebo. Anti-IL-1β treatment in CANTOS was associated with significantly lower total cancer mortality versus placebo leading to prospective clinical trials in lung cancer including trials combining anti-IL-1β and anti-PD-1 (NCT03631199).(*41, 42*) Building on this collective work, our findings raise the hypothesis that targeting the NLRP3 inflammasome or IL-1β may reverse the inflammatory phenotype of low 2IR score monocytes/macrophages and may represent a rational combination strategy to overcome CPI resistance in a defined subset of patients with UC. However, the effects of targeting the inflammasome and specific cytokines may be time and context dependent and additional preclinical and clinical work is warranted to refine this hypothesis.

Other studies exploring human specimens, often focused on individual parameters, have linked aspects of tumor-associated inflammation and/or myeloid cells with resistance to CPI in UC and other malignancies. Features reflecting an activated tumor stroma, including EMT- and TGFβ-related gene signatures have been correlated with poor outcomes with CPI treatment.(*7, 20*) Quantitative assessment of myeloid-derived suppressor cells and/or specific cytokines/chemokines in the peripheral blood have also been correlated with CPI resistance.(*43–45*) Recently, elevated levels of IL-8 in peripheral blood and in tumors, traced primarily to the myeloid cell compartment, was associated with decreased efficacy of CPI across several tumor types.(*22, 46*) Our results confirm and extend these findings providing a holistic view of the relative contribution of various components of the UC TME to CPI outcomes and highlighting the importance of assessing the balance of measures of adaptive immunity versus tumor-associated inflammation in individual tumors and cell populations.

There are potential limitations to our study. Given the paucity of human scRNA-seq data in UC to date, our scRNA-seq dataset contributes to an understanding of the cellular diversity within the UC TME but a larger cohort is required to establish a definitive cellular atlas of UC specimens. Still, the main goal of our scRNA-seq cohort in this study was to uncover the origins our gene modules derived from bulk RNA sequencing data. Features of urothelial cancer cells are likely associated with sensitivity and resistance to CPI. However, beyond TMB, on which our three gene modules were conditioned, our current analysis was focused on the TME given the expression of the module genes when projected onto our scRNA-seq data. Cancer cell-intrinsic features that contribute to immune escape and ultimately shape the “chronically” inflamed TME require further study. Together, these considerations underscore the need for additional studies of UC specimens profiled at single cell resolution and linked to CPI treatment outcomes.

Our study identified three key gene modules associated with sensitivity or resistance to CPI in patients with metastatic UC related to the balance of adaptive immunity versus tumor-associated inflammation, defined the 2IR score as reflecting such balance in individual UC TMEs and single monocytes/macrophages, and identified putative therapeutic targets to overcome resistance associated with proinflammatory monocytes/macrophages. Future work integrating the 2IR score into clinical trials seeking to overcome monocyte/macrophage-related CPI resistance, further defining “master regulators” of the low 2IR score monocyte/macrophage phenotype, and dissecting the dominant mechanisms of immune suppression related to these cells may foster precision immunotherapy in UC.

## Materials and Methods

### Identification and validation of gene modules associated with CPI outcomes from bulk RNA sequencing data

#### Patient cohorts with tumor mutational burden data and/or bulk RNA sequencing data

Three datasets including bulk RNA sequencing (RNAseq) data from patients with urothelial cancer were analyzed in this study (Figure 1 and Supplemental Table S1): IMvigor 210, The Cancer Genome Atlas (TCGA) urothelial bladder cancer dataset, and the Checkmate 275 study.

IMvigor 210 was a single arm phase 2 study investigating PD-L1 inhibition with atezolizumab (1200 mg intravenously every 3 weeks) in patients with metastatic urothelial cancer (NCT01208652, NCT02951767). The primary endpoint of the trial was the objective response rate according to Response Evaluation Criteria In Solid Tumors v1.1. Patients with metastatic urothelial cancer progressing despite prior platinum-based chemotherapy, or chemotherapy-naïve patients who were not eligible for cisplatin-based chemotherapy, were eligible. The results of IMvigor 210 have previously been reported.(*1, 14*) Patients enrolled on IMvigor 210 were required to have archival tumor tissue obtained within two years of study entry submitted for analysis which including bulk RNAseq as well as targeted next-generation sequencing-based genomic profiling for 395 cancer-related genes (FoundationOne, Foundation Medicine, Cambridge MA). These analyses were previously reported in detail by the investigators.(*7*) For the current study, RNAseq data, TMB (“FMOne mutation burden per MB”), objective response rate, and survival data for 348 unique patients were extracted from the R package *IMvigor210CoreBiologies* (http://research-pub.gene.com/IMvigor210CoreBiologies/)

The Cancer Genome Atlas bladder cancer dataset includes patients with clinically localized muscle-invasive urothelial cancer of the bladder who underwent radical cystectomy. This cohort has previously been described in detail(*47*) and RNAseq data (“Level_3_RSEM_genes_normalized”) for 408 unique patients was downloaded from Firehose (2016_01_28) at the Broad Institute (https://confluence.broadinstitute.org/display/GDAC/Home/). The updated clinical data were downloaded from an integrated TCGA pan-cancer clinical data resource.(*48*)

Checkmate 275 was a single arm phase 2 study investigating PD-1 inhibition with nivolumab (3 mg/kg intravenously every 3 weeks) in patients with metastatic urothelial cancer (NCT02387996). The primary endpoint of the trial was the objective response rate according to Response Evaluation Criteria In Solid Tumors v1.1. Patients with metastatic urothelial cancer progressing despite prior platinum-based chemotherapy were eligible. The results of Checkmate 275 have previously been reported.(*3*) Patients enrolled on Checkmate 275 were required to have archival tumor tissue submitted for analysis which including bulk RNAseq and whole exome sequencing. Patients who were consented for genomic studies and had tumor material that passed quality control were included in the current analysis. RNAseq and tumor mutational burden data was provided by Bristol Myers-Squibb and the latter was calculated as the missense mutation count. TMB (n=139) and/or RNAseq (n=72) data, objective response rate, and survival data was available with both TMB and RNAseq data available for 54 patients.

#### Preprocessing of bulk RNAseq expression datasets

For the IMvigor 210 dataset, only genes with a read count >1 in more than 10% of the samples were considered. The raw read count data from the IMvigor and Checkmate 275 datasets were first transformed to RPKM and then scaled patient-wise such that the 75% quantile of each sample was equal to 1000 (similar to the RSEM normalization(*49*)). To facilitate analysis across datasets, only 16339 genes common among the three datasets were analyzed in this study. Batch effects were removed across the three datasets using R package *ComBat*.(*50*)

#### Step-wise identification of consistently co-expressed gene modules (CCGMs)

With the goal of identifying consistently coexpressed gene modules associated with survival, we first identified genes nominally associated with better overall survival outcomes in the IMvigor 210 dataset. A bivariable Cox regression model was used to estimate the association between the expression of each gene, *Gene_i_*, with the overall survival conditional on TMB: *Surv*(*Event, Time*) ∼ *Gene_i_* + log(*TMB*). We identified 1193 genes for which higher expression was associated with better survival outcomes (nominal P-value of two-sided Wald’s test <0.05). We employed a lenient P-value cutoff to be as inclusive as possible at this initial gene selection step, and then identified <u>c</u>onsistently <u>c</u>o-expressed <u>g</u>ene <u>m</u>odules (CCGMs) to enrich for true signals and filter out possible noise. A CCGM is defined as a list of genes that are co-regulated in multiple datasets. Using weighted correlation network analysis(*51*) among these 1193 genes, we identified one co-expression module in the IMvigor 210 dataset (735 genes) and one co-expression module in the TCGA dataset (600 genes). Significant overlap (575 genes) existed between the modules identified in the IMvigor 210 and TCGA datasets (p<1e-16 by two-sided Fisher’s exact test) and these 575 overlapping genes were considered a CCGM, referred to as the “adaptive_immune_response” module

Next, we identified genes associated with worse survival outcomes conditioned on both TMB and the adaptive_immune_response module genes (*M_adaptive_immune_*). Specifically, we assessed the association of each gene (*Gene_i_*) with overall survival using a multivariable Cox regression model *Surv*(*Event, Time*) ∼ *Gene_i_* + *M_adaptive_immune_* + log (*TMB*)), where *M_adaptive_immune_* was calculated for each sample by averaging expression of the adaptive_immune_response module genes. A total of 1498 genes were associated with worse survival outcomes (nominal P-value of two-sided Wald’s test <0.05). The weighted correlation network analysis was conducted for the 1498 genes in both the IMvigor 210 and TCGA datasets, followed by the overlapping analysis analogous to the methodology described for derivation of the adaptive_immune_response module which resulted in two CCGMs, i.e. the “inflammatory_response” module (437 genes) and stromal module (287 genes). A third CCGM (50 genes) enriched with HALLMARK_MYC_targets was not further pursued in this study given its small size. We further updated the “adaptive_immune_response” module by excluding genes associated with worse survival in this three variate Cox regression model (Z>1.5), resulting in 483 genes in the module.

To investigate the pathways enriched in each CCGM, we compared the module genes with the HALLMARK and canonical gene sets in the Molecular Signatures Database (MSigDB, software.broadinstitute.org/gsea/msigdb)(*52*) using Fisher’s exact test (nominal p-value of two-sided test <1e-5).

#### Calculation of 2IR score

The expression of the adaptive_immune_response (*M_adaptive_immune_*), inflammatory response (*M_inflammatory_*), and stromal module for each sample was calculated by averaging the expression level of corresponding module genes (in the log-scale). The 2IR score was calculated as 2*IR* = *M_adaptive_immune_* − *w* × *M_inflammatory_*. The weight *w* was estimated by the relative coefficients in the Cox regression model using the IMvigor 210 training dataset: *Surv*(*Event, Time*) ∼ *w*_1_ × *M_adaptive_immune_* – w_2_ × *M_inflammatory_* and *_w_* = *w*_2/_*w*_1._ Thus, *w* represents the relative contribution of *M_inflammatory_* and *M_adaptive_immune_* to the survival outcome. Interestingly, *w* can also be interpreted as accounting for the partial variation of *M_adaptive_immune_* explained by *M_inflammatory_* score. Particularly, modeling the relationship between *M_adaptive_immune_* and *M_inflammatory_* using linear regression *M_adaptive_immune ∼ w* ×_ M_inflammatory_* the estimated *w*\* was very close to *w*_2_⁄*w*_1_. Using IMvigor 210 dataset, *w* was estimated to be 0.86 and *w*^∗^ was estimated to be 0.84. The same *w* learned from IMvigor210 dataset was applied to the validation dataset (Checkmate 275)..

#### Identifying the top-ranked genes in each module

To select a shorter list of genes in the adaptive_immune_response and inflammatory_response module with equivalent predictive power, we calculated module expression using average expression of the top N genes in each module (ranked based on their individual association with OS), from which the 2IR score was then calculated (noted as 2IR_topN). To select the optimal number of genes from each module, we plotted the association of 2IR_topN with OS against N, where N ranged from the top 1 gene in the list to the top 200 genes in the module. We then selected the “elbow point” of the curve as the optimal number of top-ranked genes for each module (Figure S4). This resulted in 10, and 39 genes in the adaptive_immune_response and inflammatory_response, respectively. Similarly, we selected 25 top genes in the stromal module by evaluating the association of adaptive_immune_response VS stromal module expression ratio with OS.

#### Univariable and multivariable models

Cox proportional hazard regression models (*coxph*() function) were performed using the R package *survival* to evaluate the association between the gene modules and TMB with OS. When module expression and TMB were treated as continuous variables, they were standardized to N(0, 1) before entering the Cox regression model to estimate hazard ratio and confidence interval, and the significance testing was performed by Wald’s test. When the 2IR score was discretized into tertiles, the R package *survminor* was used to plot the Kaplan Meier curve, and the significance testing for differences in OS was performed using the log-rank test. Logistic regression models were performed to evaluate the association between the gene modules and TMB with objective response. In the logistic regression, a complete response or partial response were treated as 1, and stable disease or progressive disease were treated as 0. The module expression and TMB were similarly standardized before entering the logistic regression model to estimate the coefficient, and the significance testing was performed by Wald’s test. All statistical analyses and figures were generated in R version 3.6.3.

#### Comparison with other biomarkers

The TIDE score was calculated using the web tool (http://tide.dfci.harvard.edu/). The CD8 effector score was calculated by averaging the CD8 effector genes as previously described.(*7*) To derive a combinatory score incorporating both the CD8 effector score and TGFB1 expression, we used a similar strategy to the calculation of 2IR score. Specifically, we used the bivariate cox regression model *Surv*(*Event, Time*) ∼ *TGFB*1 + *CD*8*effector* to learn the optimal weight for the two variables in IMvigor 210 dataset, and applied the same weight to the validation dataset of Checkmate275. A similar strategy was used to combine CXCL13 expression and ARID1A mutation.

### Multiplex immunohistochemistry and tissue analysis

#### Immunohistochemistry

For immunohistochemical staining of a subset of specimens from the Checkmate 275 cohort, we employed a tissue profiling approach known as multiplexed immunohistochemical consecutive staining on a single slide as we have previously described.(*24*) Briefly, 4-μm thick FFPE sections were pre-baked in 60 °C overnight. Slides were dewaxed manually in xylene. Slides were then loaded onto automated staining platform (Bond RX, Leica Biosystems) and covered with covertiles (Bond Universal Covertiles, Leica Biosystems) for the automatic stainer to inject the required reagents on the tissue. Peroxide block (Bond Polymer Refine Detection Kit [DS9800], Leica Biosystems) was applied for blocking the tissue endogenous peroxidase activity for 15 mins. Slides then incubated with serum-free protein block (Agilent X090930-2) for 30 mins as an extra blocking step in order to prevent nonspecific antibody binding. After the first staining cycle, Fab fragments (AffiniPure Fab Fragment Donkey anti-mouse (715-007-003) or anti-rabbit IgG (711-007-003)) against that primary antibody species were used for blocking any carry-over staining from previous immunostaining cycles using the same species of primary antibody whenever there was a repeat of same primary antibody species. Primary antibody was incubated for a certain period of time depending on its optimized protocol and a polymer detection system (Bond Polymer Refine Detection Kit [DS9800], Leica Biosystems) was used afterwards for the secondary antibody and horse radish peroxidase (HRP) binding. Slides were incubated with AEC (ImmPACT AEC Peroxide Substrate Kit, Vector Laboratories) for a specified duration defined in optimized in-house protocols for each marker. The immunostaining process was performed by an automatic immunostainer (L Bond RX, Leica Biosystems). Slides were withdrawn from the immunostainer after the automated immunostaining and counterstaining was performed manually using Modified Harris Hematoxylin (Sigma-Aldrich HHS16). Slides were then briefly incubated with ammonia water solution for crisper nuclear details and bluing purposes. Slides were mounted with Glycergel (Agilent C056330-2) and air-dried overnight. Air-dried slides were scanned by a slide scanner (NanoZoomer S60, Hamamatsu, Japan) and whole slide images were generated and stored on a server. As the preparation step for the next staining cycle on the same slides, coverslips were removed by placing the slides in a rack and immersed in hot tap water (56°C) until the mounting media dissolved. Chemical destaining was performed by immersing the slides in HCl (1%) and gradually diluted EtOH solutions after coverslip removal. The immunostaining methodology described here were repeated for all markers in the panel including PD-L1 (1:100, E1L3N, Cell Signaling), CD3 (RTU, 2GV6, Ventana), CD8 (1:100, C8/144B, Dako), CD68 (1:100, KP1, Dako), Fibronectin (1:100, F1, Abcam), and PanCK (1:50, AE1/AE3, Dako).

#### Image analysis

We imported whole slide images generated by MICSSS staining protocol(*24, 25*) into a project in the image analysis software, QuPath 0.2.0(*53*), and metadata was generated for each individual whole slide image. Accurate color vectors of hematoxylin, chromogen color (AEC) and residual color were estimated for each image by selecting a representative immunostaining zone having nearly equal amounts of all vectors. Whole section and 4 regions of interest (ROI) were annotated for the analysis. ROIs were separated into tumor and stroma by using a superpixel segmentation tool and random forest training of segmented parts. Watershed cell segmentation in combination with gaussian smoothing, intensity-based thresholding based on hematoxylin color or optical density, and diameter based cytoplasmic segmentation expansion were used for the cell segmentation. More than 30 features including minimum, maximum, mean, and standard deviation values of intensity and shape-based features were calculated and recorded during the segmentation on nucleus, cytoplasm, and total cellular compartmental area for each cell. Positive cell detection for CD8 was done by training a random forest algorithm. Intensity-based features were used with this algorithm and training was done by marking a set of segmented cells as positive or negative and feeding this training data into the random forest algorithm for the automatic classification of remainder of the segmented cells.

### Single-cell RNA Sequencing of Urothelial Cancer Specimens

#### Sample collection and specimen processing

Primary urothelial bladder cancer tumor tissue was obtained after obtaining informed consent in the context of an institutional review board approved genitourinary cancer clinical database and specimen collection protocol (IRB #10-1180) at the Tisch Cancer Institute, Icahn School of Medicine at Mount Sinai. Patients undergoing transurethral resection of bladder tumor had a portion of their tumor placed immediately into media containing 10% DMSO and 90% FBS in the operating room. The specimen was then transferred to the laboratory for further processing. Patients undergoing radical cystectomy and lymph node dissection had their bladder and lymph nodes sent directly to the pathology suite upon completion of the lymph node dissection. The bladder was bivalved and a portion of visible tumor was then placed in media as above. Adjacent normal tissue was identified in a subset of specimens based on visual inspection. The specimen was then transferred to the laboratory for further processing.

Tissue specimens were processed immediately upon receipt and dissociated into single cell suspensions using the GentleMACS Octodissociator with kit matched to the tissue type (Miltenyi Biotech) following the manufacturer’s instructions. Single-cell RNA sequencing was performed on these samples using the Chromium platform (10x Genomics, Pleasanton, CA) with the 3’ gene expression (3’ GEX) V3 kit, using an input of ∼10,000 cells. Briefly, Gel-Bead in Emulsions (GEMs) were generated on the sample chip in the Chromium controller. Barcoded cDNA was extracted from the GEMs by Post-GEM RT-cleanup and amplified for 12 cycles. Amplified cDNA was fragmented and subjected to end-repair, poly A-tailing, adapter ligation, and 10X-specific sample indexing following the manufacturer’s protocol. Libraries were quantified using Bioanalyzer (Agilent) and QuBit (Thermofisher) analysis. Libraries were sequenced in paired end mode on a NovaSeq instrument (Illumina, San Diego, CA) targeting a depth of 50,000-100,000 reads per cell. Sequencing data was aligned and quantified using the Cell Ranger Single-Cell Software Suite (version 3.0, 10x Genomics) against the provided GRCh38 human reference genome.

#### Preprocessing

Seurat(*26*) (version 3.0) was used to process the single-cell RNA sequencing data. After filtering cells with a high percentage (>20%) of mitochondrial reads and cells with >200 or <6000 genes detected, as well as potential doublets uncovered during subsequent analysis steps, 19,708 cells from 10 samples and 22,175 genes with nonzero read counts in > 5 cells were included for further analysis.

#### Identification of major cell clusters

After the read count data was log-normalized, the most variable 2000 genes were selected. Then the effect of the unique molecular identifier (UMI) count and percentage of mitochondria per cell was regressed out, followed by dimensionality reduction using principle component analysis (PCA). Finally, the cells were clustered using the *K*-nearest neighbors graph-based methods as implemented in *Seurat* (with the top 20 PC and resolution = 0.5). Cells were grouped into 9 major cell clusters based on the canonical cell-type-specific markers: T/NK (“CD3E”), B/plasma (“MS4A1”,“MZB1”,“CD79A”), DC (“HLA-DQA1”, “HLA-DQB1”), Mast (“MS4A2”), Macrophage/Monocyte (“C1QA”,“LYZ”), Endothelial (“PLVAP”), Fibroblast-related (“DCN”, “ACTA2”), Epithelial (“KRT19”) and Neuronal cells (NNAT).

#### Identification of minor cell clusters within major cell clusters

For each of the major cell clusters identified (except Mast and Neuronal cells), we further clustered cells into minor cell clusters within each major cluster. To achieve this, we applied similar steps described above to cells within each major cell cluster separately. For example, we identified the most variable 2000 genes within the T/NK cell cluster. These genes differ from the most variable genes across all cells and better reflect the variation among different subsets of T/NK cells. We then scaled the 2000 genes by removing the effect of UMI count and percentage of mitochondria per cell, based on which the top 10 PC were calculated. We used the top 10 PC instead of the top 20 PC in the minor cell clustering given that less gene variation is expected compared with major cell clustering. An additional batch effect removal step was included in the minor cell clustering. Removal of batch effect was not needed for major cell clustering as the variation among major cell types was sufficiently large to overcome the technical variation or batch effects among samples. That is, cells clustered according to major cell type rather than sample. In contrast, batch effect became more prominent in minor cell clustering. A challenge in batch effect correction is that technical artifact may be confounded by genuine sample-specific variation. This is particularly relevant for cancer cells as prior studies have shown that epithelial cancer cells were heterogeneous across different samples and grouped largely by sample versus immune and stromal cells which grouped mostly according to cell type.(*54*) Therefore, automatic batch correlation algorithms were not employed given concern for over-correlation of genuine sample-specific variation. Instead, we manually inspected the 10 PCs for each major cell cluster and removed the PCs driven mostly by sample-specific variation for non-epithelial cells leading to removal of one PC for the T/NK and one PC for the myeloid cell cluster, respectively. The cell clustering was then conducted on the remaining PCs.

To annotate each minor cell subset, marker genes were identified for each cell subset using the Seurat function *FindAllMarkers*(). Each cell subset was inspected for specific markers identified in prior analyses and annotated accordingly. A few minor cell subsets failed to show obvious subset-specific markers identified except for ribosomal and/or mitochondrial genes. The number of genes detected per cell in these subsets (median=678) was much lower compared with other subsets (median=1490). The genes in these subsets were sufficient to detect major cell cluster identity but insufficient to unambiguously determine minor cell identity. These cells (1172, 6% of total) were thus retained in the major cell clusters but removed from the minor cell clusters. Other minor cell subsets excluded from the analysis set (both major and minor cell clusters) included those expressing markers from two different major cell clusters (potential doublets, 425 cells). In total, we identified 50 minor cell clusters comprising 18,536 cells.

#### Association between cell subsets and adaptive_immune_response, inflammatory_response, and stromal modules

For each of the 9 major cell clusters, we identified cell type-overexpressed genes using *FindAllMarker()* function in *Seurat* package (with default parameters). The overlap between each of the adaptive_immune_response, inflammatory_response, and stromal module genes and overexpressed genes among the major cell clusters was assessed using odds ratio and p-value (two-sided Fisher’s exact test).

#### Identifying macrophage/monocytes with low, intermediate and high 2IR score

The AddModuleScore() function in *Seurat* package was used to calculate the module expression for each macrophage/monocyte cell for the adaptive_immune response (*M_adaptive_immune_*) and inflammatory response(*M_inflammatory_*), respectively. The 2IR score for each cell was then calculated as 2*IR* = *M_adaptive_immune_* − *w* × *M_inflammatory_*. Inspired by the interpretation of weight *w* in 2IR score for bulk samples, *w* was estimated by the coefficient *w*^∗^ in the linear regression *M_adaptive_immune_* ∼ *w*^∗^ × *M_inflammatory_* among macrophage/monocytes in our tissue scRNA-seq dataset. 2IR score was then discretized into tertiles to identify macrophage/monocytes with low, intermediate and high 2IR score.

#### NicheNet analysis

We employed *NicheNet* (R package)(*33*) to infer potential ligands regulating the genes up-regulated in tissue monocytes/macrophages with low 2IR score (342 genes). We focused our analysis on the 2022 curated ligand-receptor interactions in the *NicheNet* database (consisting of 537 and 510 unique ligands and receptors, respectively). As an input for *NicheNet* includes a list of potential ligands, we first derived a list of receptors expressed in tissue macrophage/monocytes. Specifically, we obtained the average gene expression profile for each of the 9 major cell clusters using our tumor scRNA-seq dataset (a 22,175 by 9 expression matrix) and defined a receptor as expressed in monocytes/macrophages if its corresponding expression value was larger than the median value of the expression matrix. We obtained 183 (out of the 510) receptors present in monocytes/macrophages in this manner. A list of 254 ligands interacting with these receptors, and also present in one of the 9 major cell clusters, were then considered as potential ligands. We then calculated ligand activity scores for each ligand according to its potential to regulate expression of the genes up-regulated in monocytes/macrophages with low 2IR score. The top 20 ligands were shown in the results.

### Analysis of Peripheral Blood Cell Single-cell RNA Sequencing Cohort

Single-cell RNA sequencing data for 10 frozen PBMC samples derived from pre-treatment peripheral blood of 5 patients with metastatic UC who achieved an objective response to treatment with atezolizumab and 5 patients with metastatic urothelial cancer who did not achieve an objective response to treatment with atezolizumab in the setting of the IMvigor 210 study were downloaded from GEO (GSE145281). We annotated the major cell types for each cell using the automatic annotation algorithm implemented in R package *SingleR* and the reference profiles obtained from Absolute Deconvolution.(*55, 56*) Monocytes (including both CD14 and CD16 monocytes) were further annotated into minor clusters using the strategy of unsupervised cell clustering similar to the above analysis of the tumor tissue scRNA-seq dataset. Unlike the tumor tissue scRNA-seq dataset, we observed substantial batch effect within the frozen PBMC scRNA-seq dataset: cells were clustered according to samples rather than cell types. To remove potential batch effect and focus on cell subtypes shared across samples, we used the standard workflow for data integration as recommended by the Seurat package where “anchors” were used to harmonize different datasets.(*26*) Monocytes were then clustered into 4 minor clusters based on the integrated data.

Monocytes with low, intermediate and high 2IR score were identified following the similar strategy used in the tumor tissue scRNA-seq analysis. Notably, the 2IR score was calculated for each cell using the gene expression value before batch correction or integration, such that the score was not affected by any integration procedures. Differentially expressed genes between monocytes with high and low 2IR score were also derived using the original expression before integration. Ligands regulating genes upregulated in low 2IR score monocytes were identified using *NicheNet* in the manner as described for the tumor tissue scRNA-seq dataset.

## Data availability

All relevant data are available from the authors and/or are included in the manuscript. For TCGA and IMvigor210 cohort (including the peripheral blood scRNA-seq cohort), the datasets are publicly available (see Methods for details). Bulk RNAseq, TMB counts, and clinical outcome data from the CheckMate275 from patients who consented to deposition will be submitted to dbGAP. Single cell RNA-seq data from the 10 UC tumor specimens will be deposited to GEO.

## Code availability

Software and codes used in the data analysis are either publicly available from the indicated references in the Methods section or available upon request to the authors.

## Acknowledgements

The results shown here are in part based upon data generated by the TCGA Research Network: http://cancergenome.nih.gov/.

We thank Jill Gregory at the Icahn School of Medicine Instructional Technology Group for assistance with graphic design.

## Funding

CA196521 (MDG, NB, WKO), NIH Loan Repayment Program (SI)

## Competing interests

MDG has served as a consultant or advisor to BioMotiv, Janssen, Dendreon, Merck, Glaxo Smith Kline, Lilly, Astellas, Genentech, Bristol-Myers Squibb, Novartis, Pfizer, EMD Serono, Astra Zeneca, Seattle Genetics, Incyte, Aileron, Dracen, Inovio, NuMab, Dragonfly Therapeutics. He has received research funding from Janssen, Dendreon, Novartis, Bristol-Myers Squibb, Merck, Astra Zeneca, Genentech/Roche.

NB has served as a consultant or advisor to Neon, Tempest, Checkpoint Sciences, Curevac, Primevax, Novartis, Array BioPharma, Roche, Avidea, Boehringer Ingelheim, Rome Therapeutics, Roswell Park, Parker Institute for Cancer Immunotherapy. She has received research funding from Novocure, Ludwg Institute, Celldex, Genentech, Oncovir, Melanoma Research Alliance, Leukemia & Lymphoma Society, NSTEM, Regeneron.

WKO has served as a consultant to Astellas, Astra Zeneca, Bayer, Janssen, Sanofi, Sema4 and TeneoBio.

SG reports consultancy and/or advisory roles for Merck, Neon Therapeutics and OncoMed and research funding from Bristol-Myers Squibb, Genentech, Immune Design, Agenus, Janssen R&D, Pfizer, Takeda, and Regeneron. S.G. is a named co-inventor on an issued patent for multiplex immunohistochemistry to characterize tumors and treatment responses. The technology is filed through Icahn School of Medicine at Mount Sinai (ISMMS) and non-exclusively licensed to Caprion. Mount Sinai has received payments associated with this technology and Dr. Gnjatic is entitled to future payments.

PS reports consultantships for Oncolytics, Jounce, BioAtla, Forty-Seven, Polaris, Marker, Codiak, ImaginAB, Hummingbird, Dragonfly, Lytix, Lava Therapeutics, Infinity and Achelois. Dr. Sharma reports ownership interests in Constellation, Oncolytics, Apricity Health, Hummingbird, Dragonfly, Lytix, Lava Therapeutics, Achelois, Jounce, Neon, BioAtla, Forty-Seven, Polaris, Marker, Codiak, and ImaginAb. PS also owns a patent licensed to Jounce.

JS has served as a speaker for Pfizer.

AH has served as a consultant or advisor to HTG Molecular.

JZ, LW, and ES are employees of SEMA4

## Author contributions

|  | LW | JS | KB | GA | AH | RS | AF | SG | AH | SI | PW | WO | PS | MW | KU | XY | ES | PS | NB | JZ | MG |
| --- | --- | --- | --- | --- | --- | --- | --- | --- | --- | --- | --- | --- | --- | --- | --- | --- | --- | --- | --- | --- | --- |
| Conception and design: | X |  |  | X |  |  |  |  |  |  |  |  |  |  |  |  |  |  | X | X | X |
| Financial support: |  |  |  |  |  |  |  |  |  |  |  |  |  |  |  |  |  |  | X | X | X |
| Administrative support: |  |  |  |  |  |  |  |  |  |  |  |  |  |  |  |  |  |  |  |  | X |
| Provision of study materials: |  | X |  |  |  |  |  |  |  |  |  |  | X | X | X |  |  |  |  |  | X |
| Collection and assembly of data: | X | X | X | X | X | X |  |  |  | X |  |  |  |  |  |  |  |  | X | X | X |
| Data analysis and interpretation | X | X | X | X |  | X | X | X | X | X | X | X |  |  |  | X | X | X | X | X | X |
| Manuscript writing: | X | X |  | X |  |  | X |  |  |  |  |  |  |  |  |  |  |  | X | X | X |
| Final approval of manuscript | X | X | X | X | X | X | X | X | X | X | X | X | X | X | X | X | X | X | X | X | X |
| Accountable for all aspects of the work: | X |  |  |  |  |  |  |  |  |  |  |  |  |  |  |  |  |  |  | X | X |

